# Genetic predisposition to hypouricemia on whole-exome sequencing analysis and its utilities in primary screening purposes

**DOI:** 10.1101/459727

**Authors:** Do Hyeon Cha, Heon Yung Gee, Raul Cachau, Jong Mun Choi, Daeui Park, Sun Ha Jee, Seungho Ryu, Kyeong Kyu Kim, Cheryl A. Winkler, Sung Kweon Cho

## Abstract

Differentiating between inherited renal hypouricemia and transient hypouricemia is challenging. Here, we aimed to describe the genetic predisposition of hypouricemia patients using whole-exome sequencing (WES) and assess the feasibility for genetic diagnosis in primary screening. WES was performed for the discovery of diagnostic markers in discovery cohorts (N=31). Two known genetic markers *SLC22A12* c.774G>A (p.Trp258*) and *SLC22A12* c.269G>A (p.Arg90His) were identified, We genotyped for the 2 *SLC22A12* SNPs among screened 50 hypouricemia subjects for the replication cohorts; 47 carried known SLC22A12 markers; three unexplained hypouricemic cases were analyzed by using WES. We used 46 healthy internal controls for the variant discovery. Four novel variants of *SLC22A12*, c.408C>A (p.Asn136Lys), c.674C>A (p.Thr225Lys), c.851G>A (p.Arg284Gln), and c.1285G>A (p.Glu429Lys), and one novel variant of *SLC2A9*, c. 376A>G (p.Met155Val), were identified. After filtering out known genes (*SLC22A12* and *SLC2A9*), the p.Arg78His variant in *ASB12* was overlapped in two unexplained conditions. This is the first attempt to investigate the effectiveness of integrating exome sequencing and genotype into the clinical care for hypouricemia and determine the value of genetic diagnostic screening for hypouricemia in the clinical setting. Screening of just two SNPs (p.Trp258* and p.Arg90His) identified 87.7% (71/81) of patients with hypouricemia. Early identification and intervention of hypouricemia is feasible using genetic screening to prevent acute kidney injury, especially for soldiers and athletics.

## Introduction

Uric acid (UA) is the final product of purine metabolism in humans ^1^. After reuptake in the renal proximal tubule, only 10% of filtered UA is eliminated in the urine ^2^. Serum UA level is determined by the balance between the rate of purine metabolism and clearance. Hypouricemia can be caused by malnutrition or genetic predisposition. The forms of genetic predisposition can be classified as a functional disability in UA synthesis and defects in the UA reabsorption system. Deficiencies of xanthine oxidase (XO), purine nucleoside phosphorylase (PNP), and 5-phosphoribosyl-pyrophosphate (PRPP) are related to the defects in UA synthesis ^3^. Inherited disorders of UA metabolism present as serious conditions such as intellectual disability or immunodeficiency. Renal hypouricemia (RHUC) is asymptomatic and its identification is difficult in primary medical practice unless they were presented as renal stone or exercised induced acute renal failure. The main cause is a defect in the UA reabsorption system. RHUC is a rare inherited disorder involving multiple renal UA transporters ^2^. Two types are currently reported: Type 1(OMIM: 220150) and Type 2 (OMIM: 612076). These genetically deleterious forms were identified as case reports of exercise-induced acute kidney injury (EIAKI), renal failure, and nephrolithiasis. Currently, diagnosis is based on hypouricemia (<2 mg/dL) and increased fractional excretion of UA (>10%) ^4^.

RHUC was first reported in Japan, and a mutation was found in the gene of a drug transporter in the renal proximal tubule ^5,6^. Recently, various ethnic groups including Israeli-Arab, Iraqi-Jewish, and Roma populations in the Czech Republic and Slovakia have reported renal hypouricemia ^7-12^ and have been added to the East Asian populations. Prevalence of hypouricemia is reported as 0.53% in Korea^13^ which is similar to data from the west part of Japan. Japanese data reported a geometric difference in its prevalence (0.579% of West Japanese and 0.191% of East Japanese) ^14^.

Differentiating between inherited and transient hypouricemia is challenging because a low level of UA reflects malnutrition status^15^. Moreover, a genetic utility for the diagnosis has not been conducted so far.

In this study, we investigated the genetic features of subjects with extremely low levels of UA using whole-exome sequencing (WES). Many exome-based studies have been able to detect the loss-of-function variants, missense variants and other types of variants due to changes in the triplet codon on particular genetic loci, especially in diseases with high heritability and low prevalence^16-18^. After the discovery of targeted SNPs, genetic diagnostic feasibility will be assessed in the different cohorts.

## MATERIALS AND METHODS

### Study participants

This study was approved by the institutional review board of the Kangbuk Samsung Hospital (IRB# KBSMC 2016-12-016). We screened the subjects in the Korean genome and epidemiology study (KoGES) – KoGES health examinee study (Urban Cohort) and KoGES twin and family study. Out of 179,318 individuals, the selected individuals exhibited no other syndromic features except hypouriemia. Especially, we focused on extreme hypouricemia without secondary causes (chronic kidney disease, hypertension, diabetes mellitus or any other metabolic diseases) and without any history of smoking and alcohol. Thirty-one (M:11, F:20) hypouricemic Korean genomic DNA samples were obtained from the National Biobank of Korea ^19^ and conducted WES for the discovery of causative variants. After analysis, 50 additional hypouricemic subjects without secondary causes were selected from the Korean Cancer Prevention Study (KCPS-II) cohort from the Severance Hospital, Seoul, Korea (IRB#4-2011-0277) ^20^. A total of 81 hypouricemic patients were recruited for this study. All patients had given informed consent before they were enrolled in the study, which was conducted according to the Declaration of Helsinki. The overall flowchart for this study is presented in **Figure 1.**

**Figure 1.**
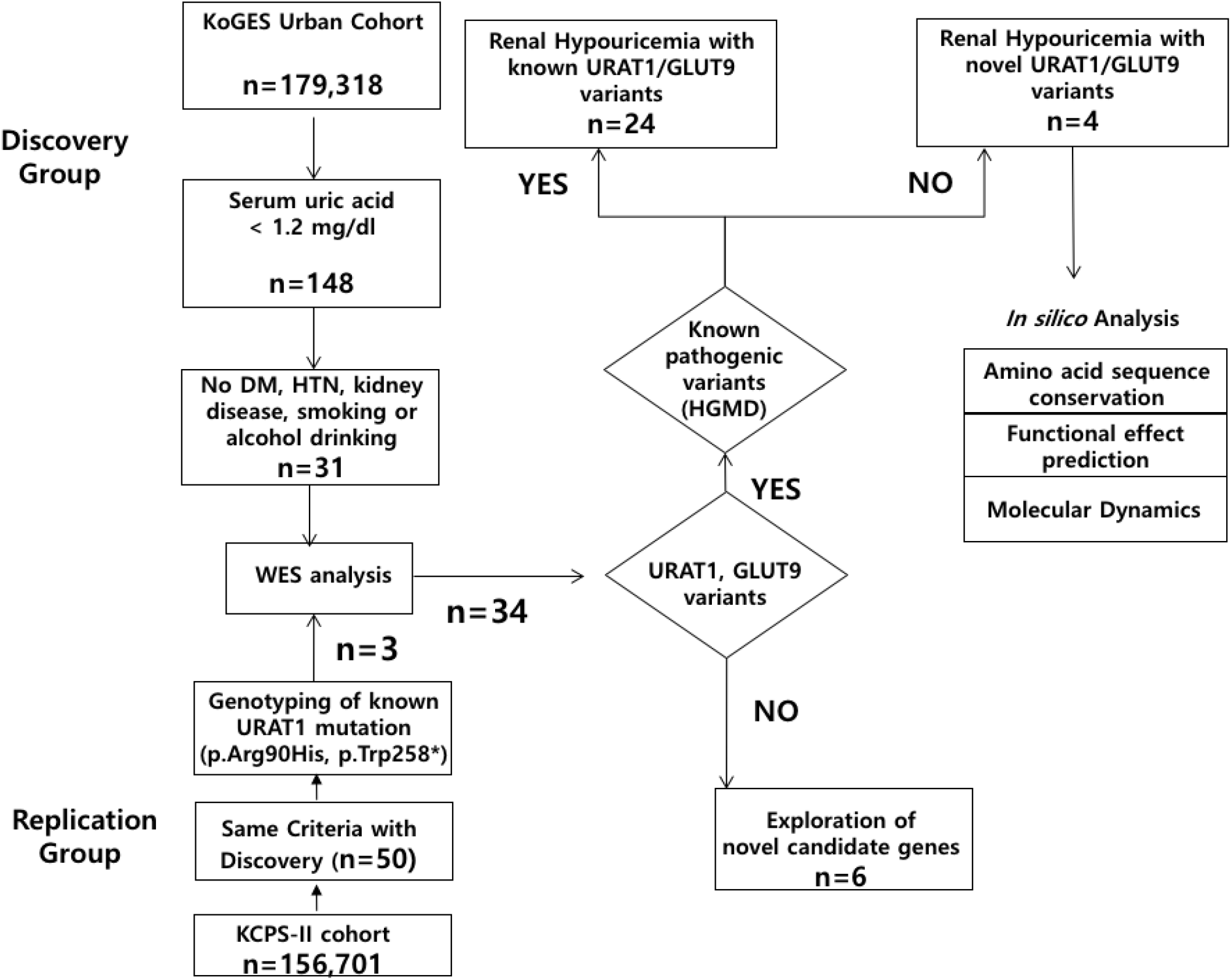
Overall flowchart for investigating novel variants associated with renal hypouricemia

### DNA preparation and whole exome sequencing

Genomic DNA was obtained from peripheral blood leukocytes. We checked the quality of the DNA with an OD260/280 ratio of 1.8–2.0 5by 1% agarose gel electrophoresis and PicoGreen^®^ dsDNA Assay (Invitrogen, Waltham, Massachusetts, USA). SureSelect sequencing libraries were prepared (Agilent SureSelect All Exon kit 50 Mb) and the enriched library was then sequenced using the HiSeq 2500 sequencing system (Illumina, San Diego, CA). Image analysis and base calling were performed with the Pipeline software using default parameters. Mapping was done by using the human reference genome assembly (GRCh37/hg19), and all variants were called and annotated using CLC Genomic Workbench (version 9.0.1) software (QIAGEN).

### WES variant filtering analysis

The overall variant-identifying process referred to the standard guidelines of investigating variants for Mendelian disorders from WES data^21,22^. We performed the analysis assuming an autosomal recessive or X-linked recessive pattern according to the observed inheritance mode in hereditary RHUC ^23^. First, based on the prevalence of hypouricemia without other medical conditions, such as hypertension or diabetes mellitus (31/179,318), the Hardy-Weinberg equation was used to calculate the allele frequency threshold to 0.01 and we excluded variants with MAF>1% in the dbSNP database (version 150), 1000 Genome Projects phase 3 data (2,504 individuals), Exome Aggregation Consortium (ExAC, http://exac.broadinstitute.org), and Genome Aggregation Database (gnomAD, http://gnomad.broadinstitute.org/)^19^. Second, variants present in the homozygous or hemizygous state in 46 healthy Koreans without hypouricemia were excluded. Third, non-synonymous variants, insertion/deletion (indel) or splice-site variants were selected. In the further analysis, we excluded single heterozygous variants so that homozygous variants and putative compound heterozygous variants finally remained. For males, hemizygous variants in the X chromosome were considered to be retained.

### Confirmation of discovered variants from WES and genotyping

Confirmation of called variants was conducted *via* direct Sanger sequencing. The DNA sequences spanning the variants were amplified using specific primers (**Supplementary Table 1**) and sequenced using an Applied Biosystems 3500xl genetic analyzer3500XL (Applied Biosystems, Foster City, CA, USA). For the screening purpose, the SNaPshot assay of rs121907896 and rs121907892 was set up (ABI PRISM SNaPshot Multiplex kit, Foster City, CA, USA) using the primer sets described in **Supplementary Table 1.**

### *In silico* analysis of novel variants

This process consists of three steps. Prior to the analysis, known pathogenic variants of *SLC22A12* and *SLC2A9* were screened in the Human Gene Mutation Database (HGMD) as a public reference. This process was performed on novel variants in *SLC22A12* and *SLC2A9*, or variants within other genes that survived the filtering analysis. The second step was to confirm that the amino acid sequences of the allelic variants were conserved across different species and was performed using the UCSC Genome Browser (https://genome.ucsc.edu/). Given the role of the nitrogen excretion function in the evolutionary process, we identified amino acid sequences in several mammals (*Rhesus macaque, Mus musculus, Canis lupus familiaris*, and *Loxodonta* a*fricana*) that share the urea cycle rather than direct UA excretion. Third, the prediction of the functional effect of variants was performed using the latest version of PolyPhen-2, SIFT, Condel, and Mutation Taster algorithms ^24-27^.

### *In silico* prediction of molecular dynamics

We initially predicted the structure of *SLC22A12* and *SLC2A9* using a homology modeling program, SWISS-MODEL (https://swissmodel.expasy.org/). The quality of predicted 3D structures was estimated on the basis of the geometrical analysis of the single model, global model quality estimation (GMQE) score and qualitative model energy analysis (QMEAN)^28^. The GenBank accession number used for each amino acid sequence was NP_653186 for *SLC22A12* and NP_064425 for *SLC2A9*. After homology modeling was completed, we selected a suitable X-ray structure for *SLC22A12* (PDB ID: 4ZW9, *SLC2A3*) and *SLC2A9* (PDB ID: 4YBQ, *SLC2A5*)^29,30^. For the more stable molecular dynamics simulations, we used I-Tasser generated models^31^. All models were generated and made publicly available and can be recovered together with the statistics from the server site (https://zhanglab.ccmb.med.umich.edu/I-TASSER/about.html). All graphical representations are made using the initial I-Tasser generated models to aid reproducibility. A qualitative evaluation of the mutation effect was evaluated based on four simple criteria. Binding urate (U) indicates the effect of the mutation on binding or urate because of the exposure of the mutated residue to the vestibular region or the urate binding motif cavity and/or involves a polar/nonpolar mutation affecting the interaction with urate. The structural effect (S) was evaluated as an increase in the root mean square displacement (RMSD) deviation computed during 25 ns of molecular dynamics (after 25 ns of equilibration) measured against the conformations obtained during a 25ns trajectory for the initial sequence using either a solvated model or a Feedback Restrained Molecular Dynamics model (FRMD). FRMD affords a simple protocol to maximally retain structural features during a molecular dynamics trajectory while minimizing distortions imposed by an external restrain^32^. The transport effect (T) indicates that the mutation intrudes into the vestibular area blocking the possible passage of urate and is assigned based on a reduction of the internal cavity volume. We used all the models to identify geometries compatible with the mutation extending the initial MD trajectory for *SLC22A12* (10 mutations) and *SLC2A9* (2 mutations) to 125 ns. All molecular dynamics calculations were performed using NAMD2^33^ and the ff99SB force field in the NVT ensemble with typical settings (T=298K, 2fs integration time, 12A cutoffs) obtained using QwikMD with default parameters to prepare the input files.

## RESULTS

### Hypouricemia prevalence and the demographic information of 81 selected hypouricemic subjects

The prevalence of extreme hypouricemia (serum UA <1.0 mg/dl) is 0.0604%, 0.032%, and 0.022% for the Kangbuk Samsung Hospital cohort (379/627,782), KoGES urban cohort (58/179,318) and KCPS-II cohort (35/156,701), respectively. After conducting WES of 31 individuals without secondary causes, we identified two potential genetic markers *SLC22A12* c.774G>A (p.Trp258*) and *SLC22A12* c.269G>A (p.Arg90His) in KoGES urban cohort. We screened an additional 50 hypouricemic individuals selected from KCPS-II cohort for the two genetic markers **(Supplementary Table 2)**. 47 individuals have known genetic markers. The genetic background of three individuals without known genetic biomarkers was further evaluated using WES. The baseline characteristics of 81 participated individuals are summarized in **Table 1.** The 81 participants (UA 0.74 ± 0.24 mg/dl; age, 47 ± 10 years; BMI, 23.3 ± 2.3 kg/m^2^; total cholesterol level, 189.2 ± 26.5 mg/dl) were healthy without any disease for participation in the present study.

**Table 1.**
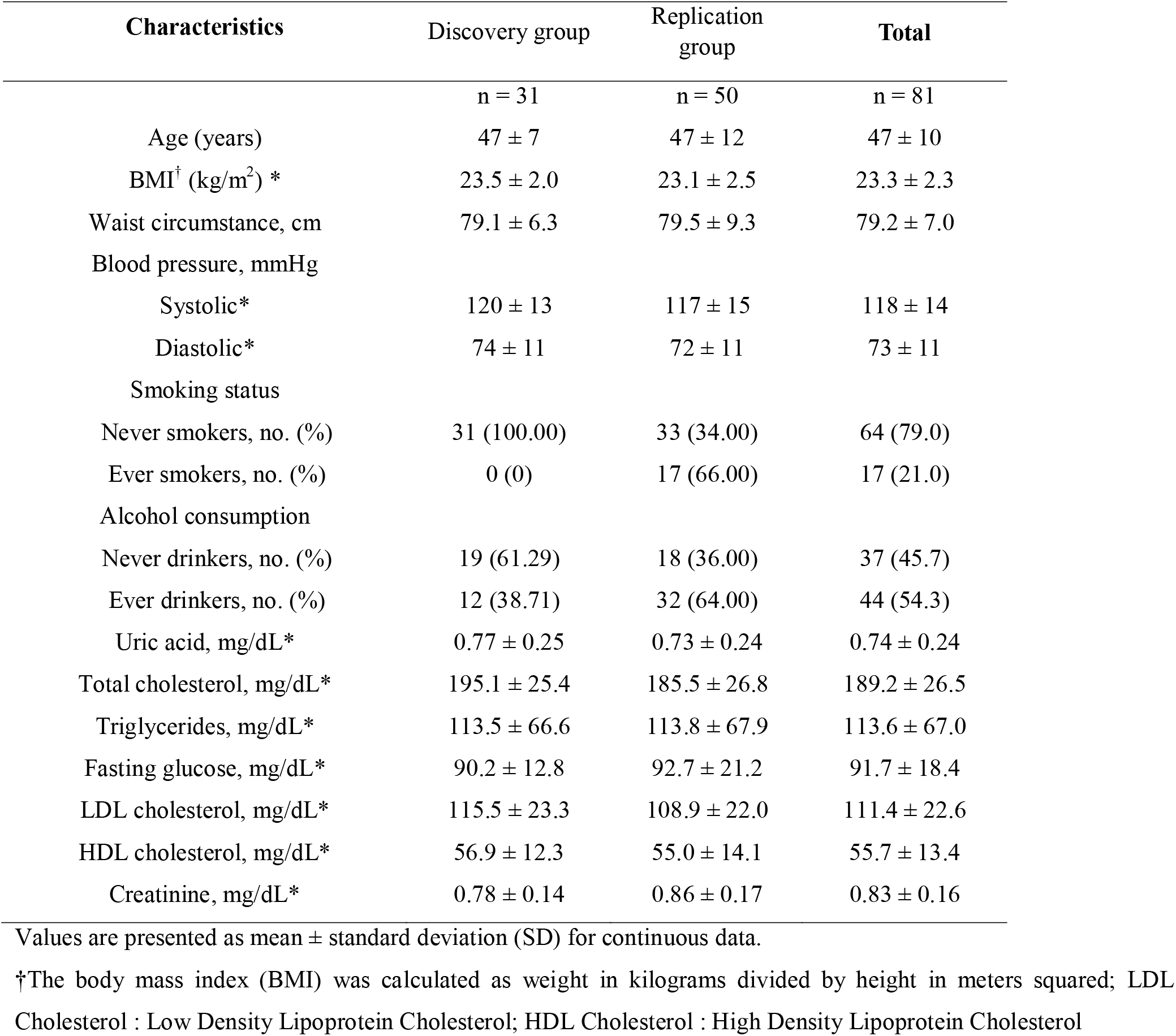
Demographic characteristics.

| Characteristics | Discovery group | Replication group | Total |
| --- | --- | --- | --- |
|  | n = 31 | n = 50 | n = 81 |
| Age (years) | 47 ± 7 | 47 ± 12 | 47 ± 10 |
| BMI<sup>†</sup> (kg/m<sup>2</sup>) * | 23.5 ± 2.0 | 23.1 ± 2.5 | 23.3 ± 2.3 |
| Waist circumference, cm | 79.1 ± 6.3 | 79.5 ± 9.3 | 79.2 ± 7.0 |
| Blood pressure, mmHg |  |  |  |
| Systolic* | 120 ± 13 | 117 ± 15 | 118 ± 14 |
| Diastolic* | 74 ± 11 | 72 ± 11 | 73 ± 11 |
| Smoking status |  |  |  |
| Never smokers, no. (%) | 31 (100.00) | 33 (34.00) | 64 (79.0) |
| Ever smokers, no. (%) | 0 (0) | 17 (66.00) | 17 (21.0) |
| Alcohol consumption |  |  |  |
| Never drinkers, no. (%) | 19 (61.29) | 18 (36.00) | 37 (45.7) |
| Ever drinkers, no. (%) | 12 (38.71) | 32 (64.00) | 44 (54.3) |
| Uric acid, mg/dL* | 0.77 ± 0.25 | 0.73 ± 0.24 | 0.74 ± 0.24 |
| Total cholesterol, mg/dL* | 195.1 ± 25.4 | 185.5 ± 26.8 | 189.2 ± 26.5 |
| Triglycerides, mg/dL* | 113.5 ± 66.6 | 113.8 ± 67.9 | 113.6 ± 67.0 |
| Fasting glucose, mg/dL* | 90.2 ± 12.8 | 92.7 ± 21.2 | 91.7 ± 18.4 |
| LDL cholesterol, mg/dL* | 115.5 ± 23.3 | 108.9 ± 22.0 | 111.4 ± 22.6 |
| HDL cholesterol, mg/dL* | 56.9 ± 12.3 | 55.0 ± 14.1 | 55.7 ± 13.4 |
| Creatinine, mg/dL* | 0.78 ± 0.14 | 0.86 ± 0.17 | 0.83 ± 0.16 |

Values are presented as mean ± standard deviation (SD) for continuous data.
†The body mass index (BMI) was calculated as weight in kilograms divided by height in meters squared; LDL Cholesterol : Low Density Lipoprotein Cholesterol; HDL Cholesterol : High Density Lipoprotein Cholesterol

### Identification of causative mutations by whole-exome sequencing

WES analysis was performed in the first cohort of 31 patients of the KoGES cohort. The average depth coverage for these individuals was 85-fold. We performed variant calling and downstream filtering analyses assuming an autosomal recessive inheritance pattern. As a result of the WES analysis, variants of *SLC22A12* and *SLC2A9* were observed in 28 of the 31 individuals, and in the remaining six individuals, variants within other genes that appeared to have possible disease-causing potential were further investigated. In the 28 people with *SLC22A12* and *SLC2A9* mutations, 24 individuals had variants previously reported in the HGMD, and the remaining four individuals had novel mutations that had not been reported previously. In the 28 subjects, we found 10 individuals with homozygous p.Trp258* variants, the founder mutation of *SLC22A12*, which is the most disease-related variant reported so far. We also identified two individuals with homozygous p.Arg90His variants ^34^. We found 12 individuals with compound heterozygous variants previously reported in the HGMD (**Table 2**). In one of the four individuals with novel variants, we found a p.Met155Val missense mutation, which was a compound heterozygous with p.Arg380Trp, a known variant of *SLC2A9* (NIH17A8568242). In another patient, the p.Glu429Lys mutation was a compound heterozygous with p.Trp258* in *SLC22A12* (NIH17A8865148). Two novel *SLC22A12* variants, the p.Thr225Lys mutation and the p.Arg284Gln mutation, were identified as compound heterozygous (NIH17A8798528). Finally, it was also confirmed that the novel p.Asn136Lys mutation was heterozygous with a previously reported p.Leu418Arg mutation (NIH17K4930892) (**Table 2**). Detailed properties of the four mutations on *SLC22A12* and one mutation on *SLC2A9* are shown in **Table 2** and **Table 3**. The overall distribution of allele frequencies of *SLC22A12* variants, which are previously reported in HGMD or are newly discovered, is shown in **Figure 2**. The novel *SLC22A12* and *SLC2A9* variants were confirmed in the participant DNA samples by direct Sanger sequencing (**Figure 3**). We used several methods for the functional prediction of the three *SLC22A12* variants and one *SLC2A9* variant found above. Amino acid sequence conservation was compared with *R. macaque, M. musculus, C. lupus familiaris*, and *L. africana* (**Table 3**). All four prediction tools (Mutation Taster, Polyphen2, SIFT, Condel) indicated that the two *SLC22A12* variants (p.Thr225Lys, p.Arg284Gln) were found to be deleterious for the function of the encoded proteins in NIH17A8798528. p.Glu429Lys of *SLC22A12* found in NIH17A8865148 was only predicted to be disease-causing in Mutation Taster, whereas it was predicted to be benign in the remaining tools. In the case of p.Asn136Lys, however, all prediction tools indicated that this amino acid change was not deleterious. No prediction result was obtained for p.Met155Val of *SLC2A9* found in NIH17A8568242 (**Table 3**).

**Figure 2.**
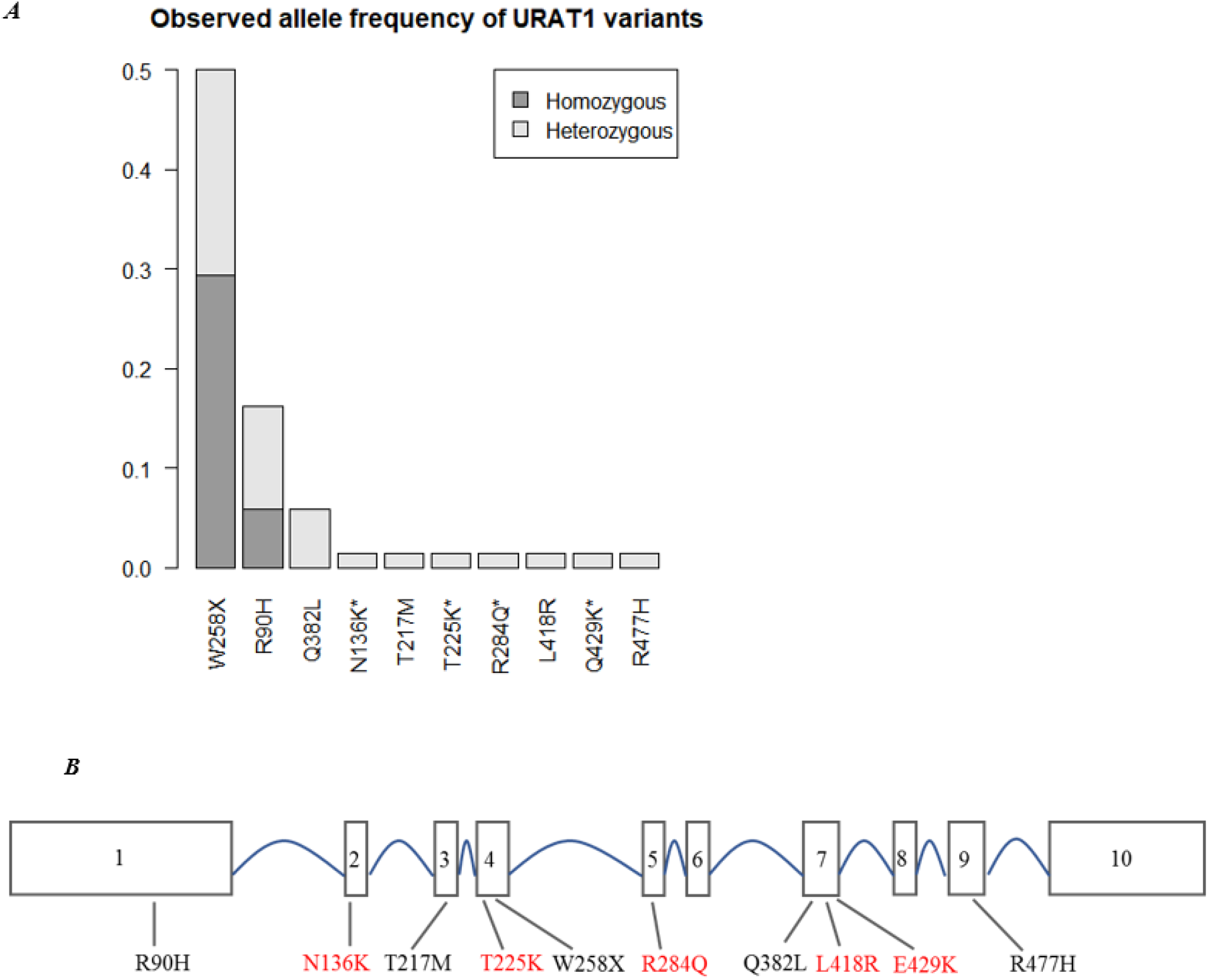
(A) Allele frequency distribution in *SLC22A12* variants. (B) A schematic diagram of the exonic location of *SLC22A12* variants found in 28 subjects. Newly discovered coding variants are marked in red

**Figure 3.**
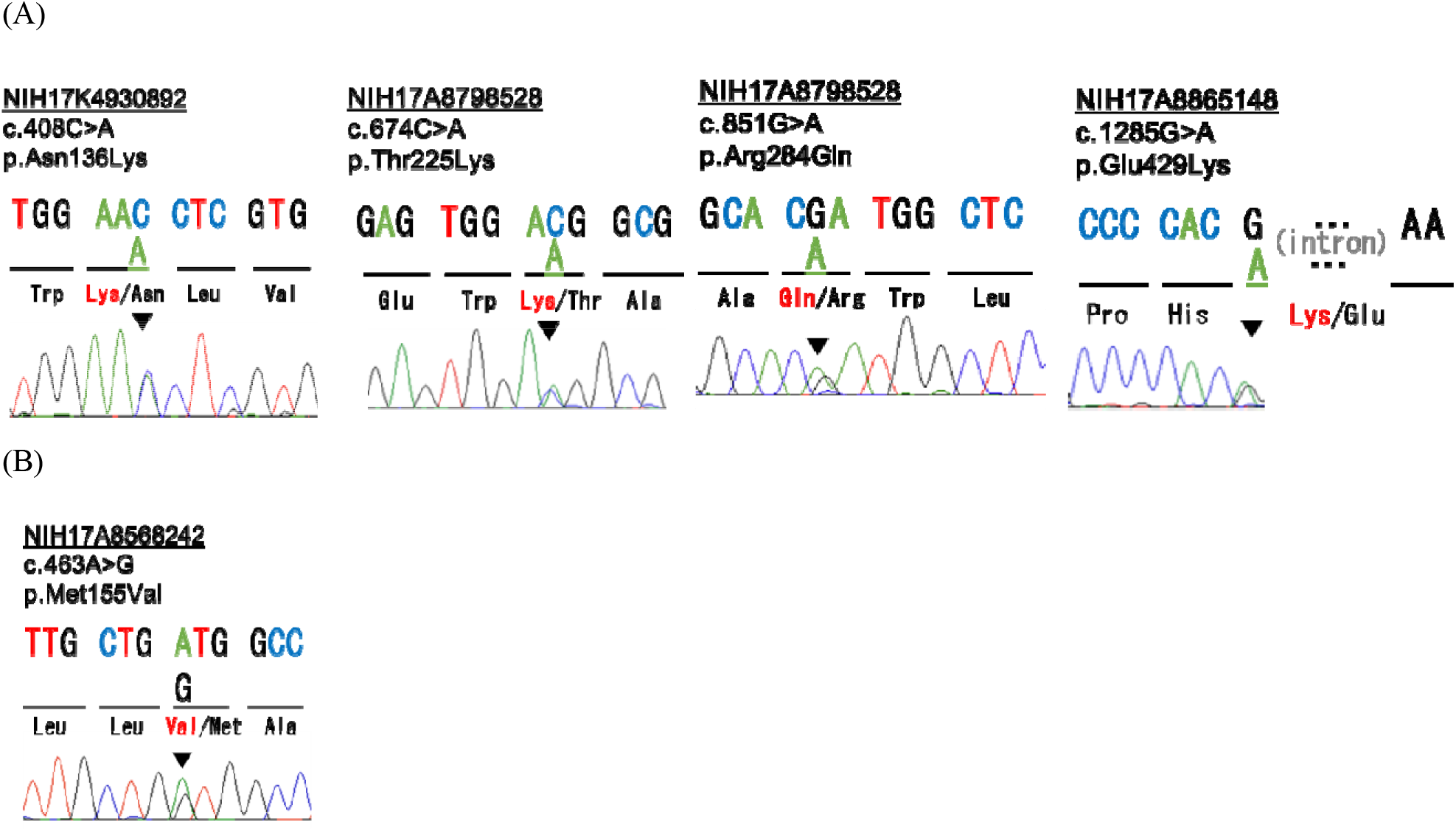
Sequencing traces of variants detected in (A) *SLC22A12* and (B) *SLC2A9.*

**Table 2.** Bi-allelic mutations of *SLC22A12* and *SLC2A9* in 28 individuals with hypouricemia.

| Individual | Gene | Zygoty | Genomic<br>change | Amino acid<br>change | Location | Mutation<br>type | HGMD® |
| --- | --- | --- | --- | --- | --- | --- | --- |
| NIH17A8087761 | *SLC22A12* | Homozygous | c.774G>A | p.Trp258* | Exon 4 | Loss-of-function | Yes |
| NIH17A8113116 | *SLC22A12* | Homozygous | c.774G>A | p.Trp258* | Exon 4 | Loss-of-function | Yes |
| NIH17A8172895 | *SLC22A12* | Homozygous | c.774G>A | p.Trp258* | Exon 4 | Loss-of-function | Yes |
| NIH17A8208690 | *SLC22A12* | Homozygous | c.774G>A | p.Trp258* | Exon 4 | Loss-of-function | Yes |
| NIH17A8214059 | *SLC22A12* | Homozygous | c.774G>A | p.Trp258* | Exon 4 | Loss-of-function | Yes |
| NIH17A8441978 | *SLC22A12* | Homozygous | c.774G>A | p.Trp258* | Exon 4 | Loss-of-function | Yes |
| NIH17A8503979 | *SLC22A12* | Homozygous | c.774G>A | p.Trp258* | Exon 4 | Loss-of-function | Yes |
| NIH17A8555604 | *SLC22A12* | Homozygous | c.774G>A | p.Trp258* | Exon 4 | Loss-of-function | Yes |
| NIH17A8900052 | *SLC22A12* | Homozygous | c.774G>A | p.Trp258* | Exon 4 | Loss-of-function | Yes |
| NIH17A8951027 | *SLC22A12* | Homozygous | c.774G>A | p.Trp258* | Exon 4 | Loss-of-function | Yes |
| NIH17A8293564 | *SLC22A12* | Homozygous | c.269G>A | p.Arg90His | Exon 1 | Missense | Yes |
| NIH17A8668884 | *SLC22A12* | Homozygous | c.269G>A | p.Arg90His | Exon 1 | Missense | Yes |
| NIH17A8265785 | *SLC22A12* | Compound | c.774G>A | p.Trp258* | Exon 4 | Loss-of-function | Yes |
|  |  | Heterozygous | c.269G>A | p.Arg90His | Exon 1 | Missense | Yes |
| NIH17A8304859 | *SLC22A12* | Compound | c.774G>A | p.Trp258* | Exon 4 | Loss-of-function | Yes |
|  |  | Heterozygous | c.269G>A | p.Arg90His | Exon 1 | Missense | Yes |
| NIH17A8429119 | *SLC22A12* | Compound | c.774G>A | p.Trp258* | Exon 4 | Loss-of-function | Yes |
|  |  | Heterozygous | c.269G>A | p.Arg90His | Exon 1 | Missense | Yes |
| NIH17A8517296 | *SLC22A12* | Compound | c.774G>A | p.Trp258* | Exon 4 | Loss-of-function | Yes |
|  |  | Heterozygous | c.269G>A | p.Arg90His | Exon 1 | Missense | Yes |
| NIH17A8595401 | *SLC22A12* | Compound | c.774G>A | p.Trp258* | Exon 4 | Loss-of-function | Yes |
|  |  | Heterozygous | c.269G>A | p.Arg90His | Exon 1 | Missense | Yes |
| NIH17A8768605 | *SLC22A12* | Compound | c.774G>A | p.Trp258* | Exon 4 | Loss-of-function | Yes |
|  |  | Heterozygous | c.269G>A | p.Arg90His | Exon 1 | Missense | Yes |
| NIH17A8775970 | *SLC22A12* | Compound | c.774G>A | p.Trp258* | Exon 4 | Loss-of-function | Yes |
|  |  | Heterozygous | c.269G>A | p.Arg90His | Exon 1 | Missense | Yes |
| NIH17A8850018 | *SLC22A12* | Compound | c.774G>A | p.Trp258* | Exon 4 | Loss-of-function | Yes |
|  |  | Heterozygous | c.650C>T | p.Thr217Met | Exon 3 | Missense | Yes |
| NIH17A8449886 | *SLC22A12* | Compound | c.774G>A | p.Trp258* | Exon 4 | Loss-of-function | Yes |
|  |  | Heterozygous | c.1145A>T | p.Gln382Leu | Exon 7 | Missense | Yes |
| NIH17A8615650 | *SLC22A12* | Compound | c.774G>A | p.Trp258* | Exon 4 | Loss-of-function | Yes |
|  |  | Heterozygous | c.1145A>T | p.Gln382Leu | Exon 7 | Missense | Yes |
| NIH17A8656939 | *SLC22A12* | Compound | c.774G>A | p.Trp258* | Exon 4 | Loss-of-function | Yes |
|  |  | Heterozygous | c.1145A>T | p.Gln382Leu | Exon 7 | Missense | Yes |
| NIH17A8304077 | *SLC22A12* | Compound | c.1145A>T | p.Gln382Leu | Exon 7 | Missense | Yes |
|  |  | Heterozygous | c.1430G>A | p.Arg477His | Exon 9 | Missense | Yes |
| NIH17A8865148 | *SLC22A12* | Compound | c.774G>A | p.Trp258* | Exon 4 | Loss-of-function | Yes |
|  |  | Heterozygous | c.1285G>A | p.Glu429Lys | Exon 7-8 | Missense | No |
| NIH17A8798528 | *SLC22A12* | Compound | c.674C>A | p.Thr225Lys | Exon 4 | Missense | No |
|  |  | Heterozygous | c.851G>A | p.Arg284Gln | Exon 5 | Missense | No |
| NIH17K4930892 | *SLC22A12* | Compound | c.408C>A | p.Asn136Lys | Exon 2 | Missense | No |
|  |  | Heterozygous | c.1253T>G | p.Leu418Arg | Exon 7 | Missense | Yes |
| NIH17A8568242 | *SLC2A9* | Compound | c.1138C>T | p.Arg380Trp | Exon 10 | Missense | Yes |
|  |  | Heterozygous | c.463A>G | p.Met155Val | Exon 5 | Missense | No |

HGMD®: The Human Gene Mutation Database

**Table 3.** Novel variants of SLC22A12 and SLC2A9 identified in individuals with hypouricemia by whole-exome sequencing.

| Gene symbol | Individual | Chr | Base position | Nucleotide change<sup>a</sup> | Amino acid change | Amino acid conservation |  |  |  | Frequency in the dbSNP database<sup>b</sup> | Frequency in the gnomAD database<sup>c</sup> | Mutation Taster<sup>e</sup> | PP2 Humvar<sup>f</sup> | SIFT<sup>g</sup> | Condel<sup>h</sup> |
| --- | --- | --- | --- | --- | --- | --- | --- | --- | --- | --- | --- | --- | --- | --- | --- |
|  |  |  |  |  |  | *Rhesus macaque* | *Mus musculus* | *Canis lupus familiaris* | *Loxodonta africana* |  |  |  |  |  |  |
| *SLC22A12* | NIH17A8865148 | 11 | 64367362 | c.1285G>A | p.Glu429Lys | Glu | Gly | Glu | Glu | rs139140123<br>A=0.00005/5 (ExAC)<br>A=0.00008/1 (GO-ESP) | 0.000044 (no hom) | DC | Bn (0.37) | Tol (0.05) | Neu (0.463) |
|  | NIH17A8798528 | 11 | 64361119 | c.674C>A | p.Thr225Lys | Thr | Thr | Thr | Thr | No | No | DC | Dam (0.998) | Del (0) | Del (0.919) |
|  | NIH17A8798528 | 11 | 64366008 | c.851G>A | p.Arg284Gln | Arg | Arg | Arg | Arg | No | 0.000019 (no hom) | DC | Dam (0.527) | Del (0.03) | Del (0.542) |
|  | NIH17K4930892 | 11 | 64360256 | c.408C>A | p.Asn136Lys | Asn | Asp | Asp | Asp | No | 0.000004 (no hom) | PM | Bn (0.345) | Del (0) | Del (0.553) |
| *SLC2A9* | NIH17A8568242 | 4 | 9987365 | c.376A>G | p.Met155Val | Met | Met | Met | - | rs369512758<br>G=0.00002/3 (ExAC)<br>G=0.0002/2 (GO-ESP) | 0.000022 (no hom) |  |  |  |  |

Abbreviations are as follows: Chr, chromosome; Bn, benign; Condel, consensus deleteriousness score of non-synonymous single nucleotide variants; Dam, damaging; DC, disease causing; Del, deleterious; Neu, neutral; PM, polymorphism; PP2, PolyPhen-2 prediction score Humvar; SIFT, sorting intolerant from tolerant; SNP, single nucleotide polymorphism; To, tolerant.
<sup>a</sup>cDNA mutations are numbered according to human cDNA reference sequence NM\_144585.2 (*SLC22A12*), NM\_001001290.1 (*SLC2A9*). <sup>b</sup>dbSNP database (http://www.ncbi.nlm.nih.gov/SNP). <sup>c</sup>gnomAD browser (http://gnomad.broadinstitute.org/). <sup>e</sup>Mutation taster (http://www.mutationtaster.org/). <sup>f</sup>PolyPhen-2 prediction score HumVar ranges from 0 to 1.0; 0 = benign, 1.0 = probably damaging (http://genetics.bwh.harvard.edu/pph2/). <sup>g</sup>SIFT (http://sift.jcvi.org/). <sup>h</sup>Condel (http://bbglab.irbbarcelona.org/famnsdb/)

### Molecular dynamic prediction of *SLC22A12* and *SLC2A9* and novel variant location

The amino acid substitutions in SLC22A12 (10 mutations) and SLC2A9 (2 mutations) were considered. The predicted functional impact of the amino acid change is illustrated in **Table 5**. Our overall organization of *SLC2A9* and *SLC22A12* is similar to the molecular dynamics approach described by Clemencon et al. ^35^. Steered dynamic simulations of urate transport were performed with mutations in *SLC22A12* and *SLC2A9* and are presented in **Figure 4**. Assessing the extent of the effect of the mutations in the S set is difficult in a qualitative analysis due to the large changes observed during the molecular dynamics trajectory. p.Arg90His, p.Thr217Met, p.Thr225Lys, p.Trp258*, and p.Leu418Arg for *SLC22A12* and p.Met155Val and p.Arg380Trp for *SLC2A9* are classified as structural effect. p.Arg284Gln and p.Arg477His for *SLC22A12* are estimated as structural effect. p.Asn136Lys and p.Gln382Leu for *SLC22A12* are regarded as binding effect. p.Arg477His for *SLC22A12* may contribute to both the lower binding of urate and blocking of the transport path.

**Figure 4.**
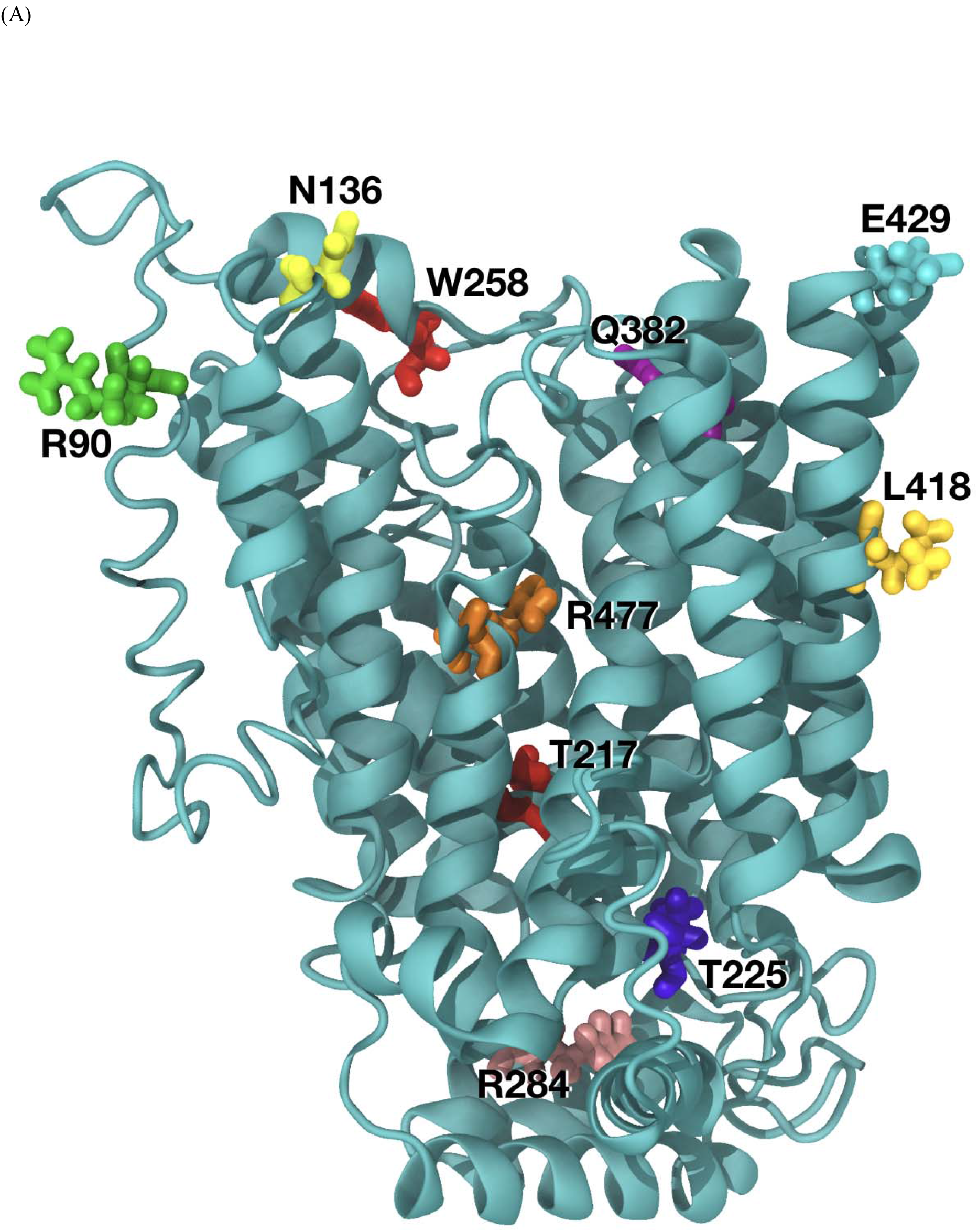
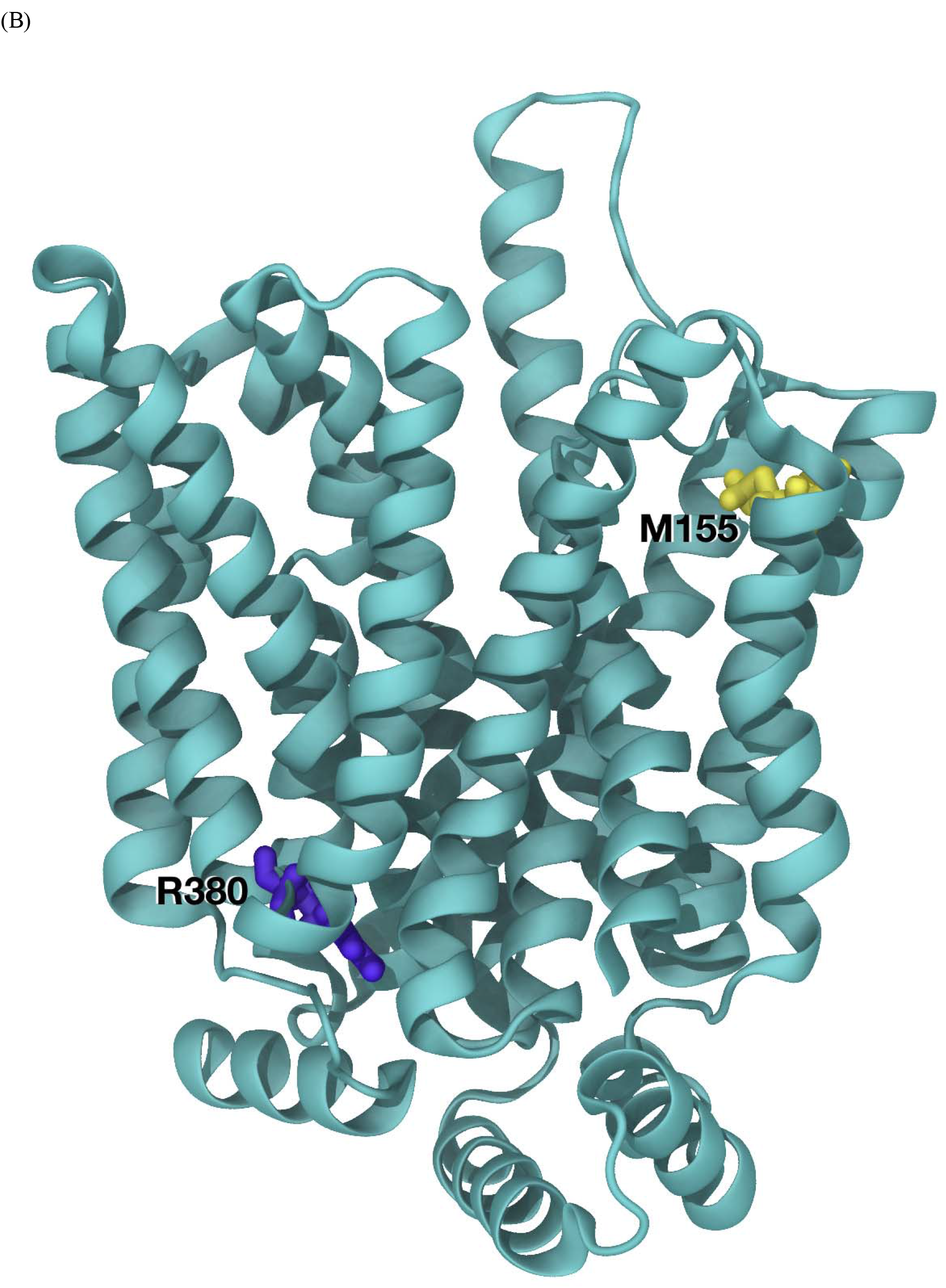
Mapping of residues to predicted model (A) *SLC22A12* Model and mutation location. (B) *SLC2A9* model and mutation location. (C) SLC2A9 model. Standard orientation top tilted 30° towards the viewer to better appreciate the disposition of residues M155 (in yellow, top left) and R380 (in yellow, bottom right). Rather unexpectedly R380 has an important structural role stabilizing the fold of the model. Its functional role is being further explored using steered dynamics.

### Screening with two genetic markers *SLC22A12* c.774G>A (p.Trp258*) and c.269G>A (p.Arg90His)

Among 50 hypouricemic individuals from the KCPS-II cohort, 47 individuals carried one of the two genetic markers; six individuals carried a c.774G>A (p.Trp258*) homozygote mutation; one individual carried a c.269G>A (p.Arg90His) homozygote mutation; 22 individuals carried a c.269G>A (p.Arg90His) and c.774G>A (p.Trp258*) compound heterozygous mutation; 15 individuals carried a c.269G>A (p.Arg90His) or c.774G>A (p.Trp258*) heterozygote mutation. The result is illustrated in **Supplementary Table 2**.

### Identification of Novel candidate genes by whole-exome sequencing

WES was conducted to resolve three cases that were not indicated by the two causative genetic markers. A total of six patients were used to investigate candidate genes associated with hypouricemia. The number of variants at each filtering step for each individual is outlined in **Supplementary Table 3**. For example, in NIH17A8004492, a total of 152,116 variants were detected from the normal human genome sequence. Of these, 18,940 variants were left after exclusion of homozygous or hemizygous variants in 46 healthy controls and the exclusion of common variants (MAF >1% in dbSNP database). After excluding the synonymous variants, the significant variants were reduced to 1,412. The detailed types of variation are described in **Supplementary Table 3**. When considering only the variants satisfying the autosomal recessive inheritance pattern, in which the MAF threshold is lower than 1% in any population group, and the amino acid sequence substitution does not occur in the reference sequence of other mammals, only 15 variants of nine genes in NIH17A8004492 had possible disease-causing potential. The variant filtering analyses were performed in the same manner for NIH17A8239849, NIH17A8738324, NIH1705180563, YID182829, and YID632847. Rare exonic variants identified through the described filtering criteria in six patients, whose WES analysis results did not contain either the *SLC22A12* or *SLC2A9* variant, are shown in **Table 4.** Overall, 9, 26 and 4 candidate genes (total of 39) were identified as homozygous mutations, compound heterozygous mutations, and hemizygous mutations, respectively. A p.Arg78His hemizygous variant (rs145118752) of Ankyrin repeats and SOCS Box-containing 12 (*ASB12)* in chromosome X was overlapped in two male individuals, NIH17A8004492 and YID182829. The detailed information of the investigated variants, including minor allele frequency in several populations and the results of *in silico* functional prediction tools, are described **Supplementary Table 4**.

**Table 4.**
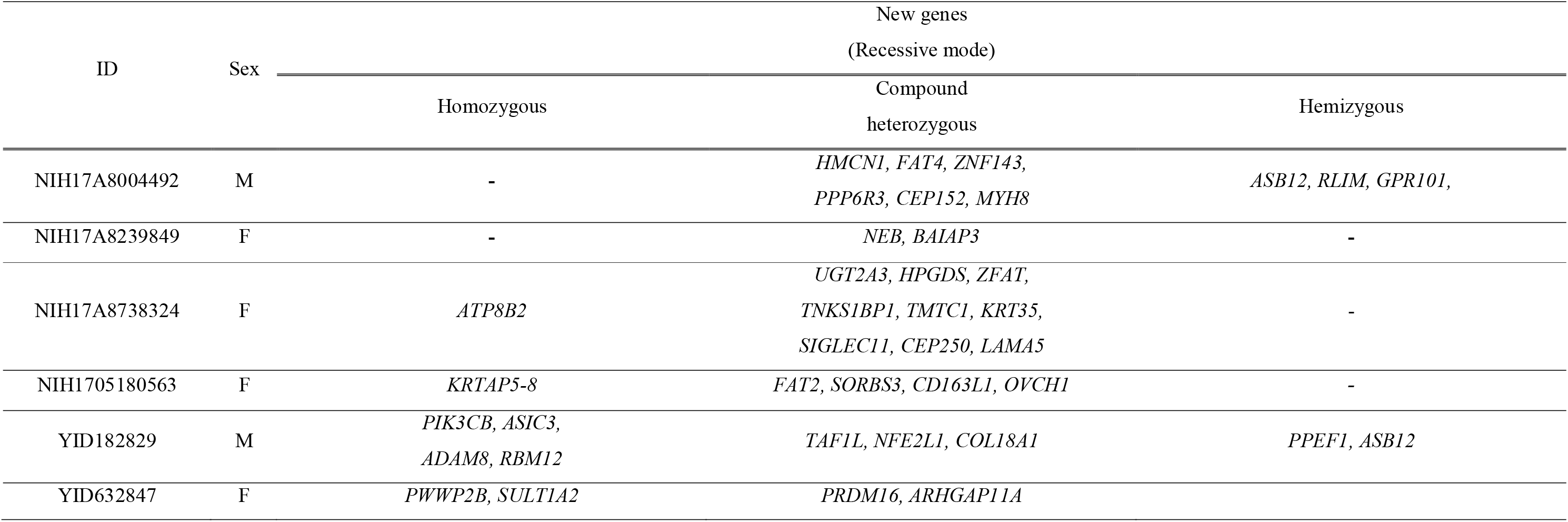
Novel genetic variants discovered in six individuals with hypouricemia.

**Table 5.**
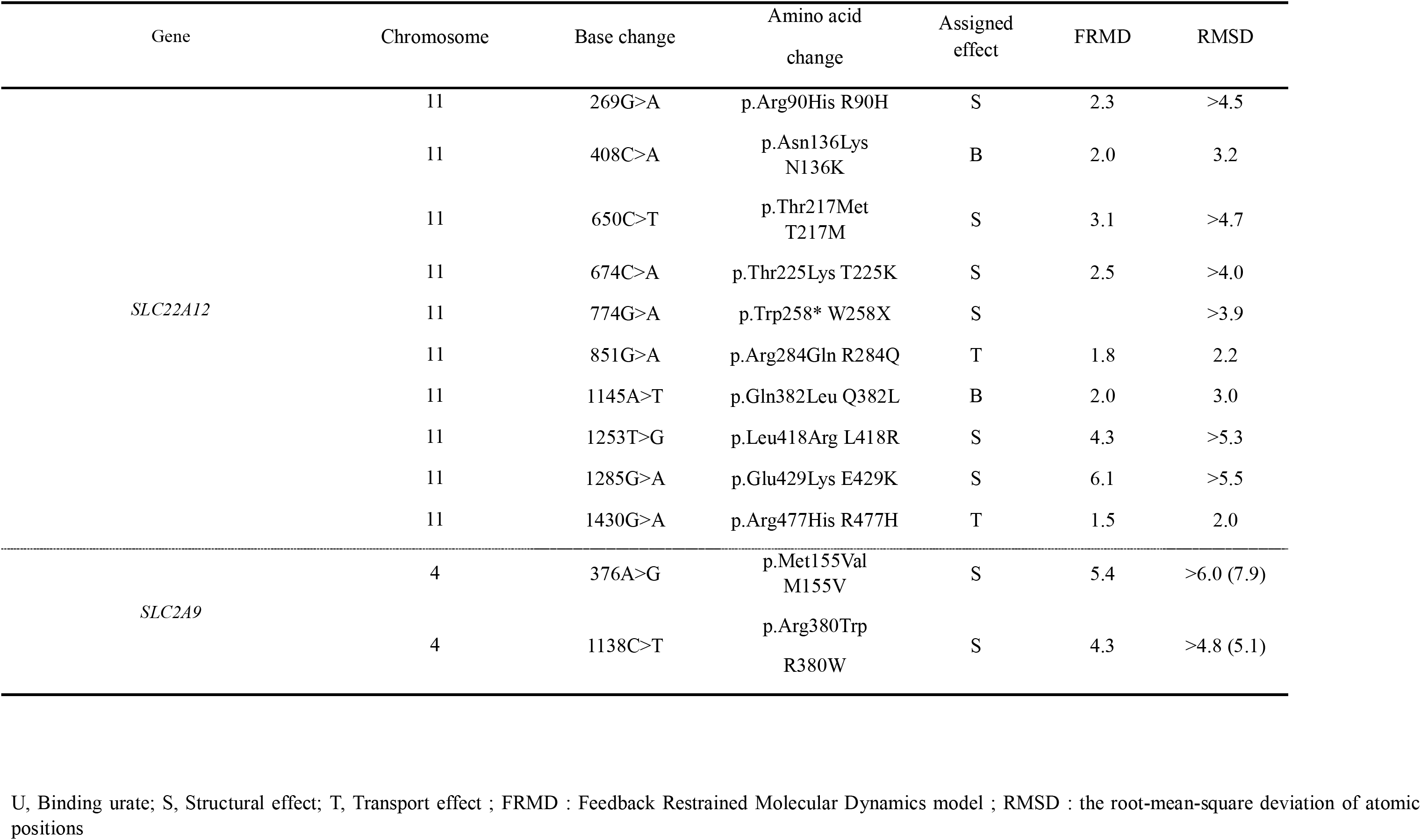
Predicted functional impact of amino acid change

## Discussion

In this study, we comprehensively evaluated the contribution of genetic predisposition to hypouricemia with WES data and genotyping date in 81 unrelated patients. Our approach is derived from that of the previous population-based studies on the adverse health effects of hypouricemia ^20,36^. Throughout our study, we were able to evaluate the genetic markers in hypouricemia which could have a potential diagnostic role for the first time. Of the 34 patients whose WES data were available, 27 had bi-allelic exonic variants within *SLC22A12* and one had a compound heterozygous variant in *SLC2A9*. We discovered four novel mutations in *SLC22A12* and one mutation in *SLC2A9* and identified the novel candidate genes in the recessive trait (Table 4).

Among the individuals with *SLC22A12* mutations, the novel variants that had not been reported before in the HGMD^®^ are p.Asn136Lys, p.Thr225Lys, p.Arg284Gln, and p.Glu429Lys. When localizing each variant on the 12 transmembrane domains of the SLC22A12 transporter, p.Asn136Lys (exon2) was located at the end of an intracellular loop, p.Thr225Lys (exon4) was present at the beginning of an extracellular loop, p.Arg284Gln (exon5) was localized in the largest extracellular loop, and p.Glu429Lys, in which the distal end of exon 7 and the first part of exon 8 are connected via splicing, was found to be within the membrane before an intracellular loop^34^. p.Asn136Lys, which was bi-allelic with p.Leu418Arg in the case of NIH17K4930892, was predicted to be deleterious in SIFT and Condel when analyzed *in silico*. p.Thr225Lys and p.Arg284Gln, two novel variants found to be compound heterozygous in NIH17A8798528, were predicted to be pathogenic in Mutation Taster, Polyphen2, SIFT, and Condel, respectively. In the case of p.Glu429Lys found with p.Trp258* in NIH17A8865148, only Mutation Taster reported that the variant could be disease-causing. Most of *SLC22A12* loss-of-function mutation (compound heterozygous and/or homozygous) are found in Japanese (OMIM #220150, RHUC1)^34,37-39^ with autosomal recessive inheritance pattern ^23^. It was reported that c.1245_1253del and c.1400C>T variants of *SLC22A12* are highly frequent in Roma population (1.87 and 5.56 %, respectively) ^40^.

With regard to *SLC2A9*, the p.Met155Val mutation was newly discovered in NIH17A8568242 (a compound heterozygote with p.Arg380Trp). Since an *in silico* prediction was not obtained, molecular dynamics supported its deleterious effect considering its RMSD value. Two exonic SNPs (p.Arg380Trp, rs121908321, and p.Arg198Cys, rs121908322) are well-known as RHUC2 with in more 10 patients as heterozygote defects^41,42^. Heterozygous mutation of *SLC2A9* has a significant effect on lowering UA level. Homozygous or compound heterozygous mutation has a much significant effect on UA lowering. UA is near 0.

*SLC2A9* is the most reported gene associated with serum UA levels along with *ABCG2* in gout and hyperuricemia^43^. Intronic SNPs (rs4529048, rs7674711, and rs11936395) of *SLC2A9* have been associated with both increased serum UA level and increased risk of gout^44,45^. Interestingly, an exonic variant (p.Val253Ile, rs16890979) was reported as a protective SNP for gout and decreasing UA level^46,47^. Moreover, *SLC2A9* showed a statistically significant gene-gene interaction with variants in the intergenic region located 80 kb downstream (*WDR1*-*ZNF518B*)^48^. A comprehensive study is needed to evaluate the effect of different transcriptional factors on the gene expression of *SLC2A9* variants and the surrounding loci. Utilizing the whole genome sequencing data in The Trans-Omics for Precision Medicine (TOPMed) program could be a next step to solve this phenomenon.

Recently, large scale WES using 19,517 participants (15,821 European Ancestry and 3,696 African Ancestry) identified that novel variants of *SLC22A12* and *SLC2A9*^49^. It is obvious that variants of lowering UA is different by ethnic group in the same genes. Collaborative international research with reported populations (i.e. Japan, China, Iraqi Jews Macedonia, the United Kingdom, and Czech Republic) is needed to further investigate new genes associated with renal hypouricemia when considering the genetic composition according to various ancestries.

For six unexplained cases, novel candidate genes were identified using the WES analysis strategy described above (**Table 4, Supplementary table 4**). In a systematic review, most of the genes are not known to be involved in biological pathways affecting UA levels. However, the p.Arg78His variant (rs145118752) of *ASB12* on chromosome X, which was discovered in both NIH17A8004492 and YID182829, is found in 0.018% of the whole population and 0.16% when limited to East Asia only (gnomAD 2018.7, http://gnomad.broadinstitute.org/). Thus, the fact that the rare variant was shared in unrelated hypouricemic individuals indicates that there is a considerable possibility of disease-causing potential in this variant. Little is known about the functional significance of this gene. Since it is postulated that the ASB family may be involved in protein degradation *via* mediating the ubiquitin-proteasome system or signal transduction^50^, it may be involved in trafficking or the intracellular degradation of the UA transporter. Further studies are needed to elucidate the association between hypouricemia and the *ASB12* variant and this is the limitation of study that six unexplained cases are not clearly explained in relation to a significant decrease in serum UA levels. Family study with unsolved cases of *SLC22A12 and SLC2A9* would be beneficial to reach more definite conclusions on the segregation patterns of variants. Given that several causative genes for hypouricemia are still unidentified, genetic inheritance of hypouricemia could be more common than indicated by our results.

The main purpose of this study was to view the composition of known and unknown genes for RHUC systematically in a population-based study. Hypouricemia is often regarded as an unrecognized or neglected disorder in a public health aspect. The prevalence of urolithiasis through excess UA excretion is 6–7 times higher in patients with RHUC than in the normal population ^34^. Reflecting the ability of UA as a powerful scavenger of peroxide radicals, accounting for up to 60% in the plasma, evidence of oxidative stress has accumulated not only in EIAKI and urolithiasis but also in neurodegenerative disease (Parkinson’s disease)^51-55^. The anti-oxidative stress hypothesis is also supported by the results of Facheris et al., which show that the SLC2A9 mutation is associated with lower serum UA and increased early onset of Parkinson’s disease ^56^. Early identification and intervention of hypouricemia (avoiding hard exercise, drinking plenty of water, and pre-emptively taking XO inhibitors) may prevent its adverse outcome in the near future, especially military personnel and athletics. XO inhibitor use (allopurinol or febuxostat) may be beneficial by lowering filtered UA. Neonatal screening or in-born screening of two SNPs (p.Trp258*/rs121907892 and p.Arg90His/rs121907896) will help in the early diagnosis of hypouricemia and increase awareness among primary care physicians or medical care professionals in army to prevent it from progressing to adverse outcomes. Since the genotyping of two SNPs costs less than 5 dollars and requires only half a day, it will be a huge advantage for medical doctors to consider this genetic test as a routine procedure. Our findings of 87.7% of patients with hypouricemia explained by two SNPs in *SLC22A12* gives strong evidence for its feasibility in diagnostic use. Our findings indicate that the genotyping of two SNPs in *SLC22A12* followed by WES is a simple and precise protocol for molecular diagnosis of sporadic cases of hypouricemia complicated by renal stones or EIAKI.

In summary, this study indicates the value of genetic screening of hypouricemia and suggests close monitoring of UA levels to prevent not only urolithiasis but also oxidative stress-induced disease progression for hypouricemia patients in the primary care setting.

## Supplements

Supplementary Table 1. Primer information for the SLC22A12 and SLC2A9 variants.

Supplementary Table 2. SNaPshot results of rs121907896 (p.Arg90His) and rs121907892 (p.Trp258*)

Supplementary Table 3. Variant filtering process of whole-exome sequencing (WES)

Supplementary Table 4. Possible variants identified in six individuals with hypouricemia by whole-exome sequencing

## Acknowledgements

The bioresources for this study were provided by the National Biobank of Korea, Centers for Disease Control and Prevention, Republic of Korea. This research was supported by the Basic Science Research Program through the National Research Foundation of Korea (NRF) funded by the Ministry of Education (NRF-2016R1A6A3A11933380 to S.K.C and 2015R1D1A1A01056685 to H.Y.G) and the Korea Health Technology R&D Project through the Korea Health Industry Development Institute (KHIDI), funded by the Ministry of Health & Welfare, Republic of Korea (Grant Number: HI17C2372 to S.K.C). We thank the Research Division, NEXBiO Co. Ltd for their contribution to our sequencing. We appreciate Young Sup Cho MD, PhD and Hyekyung Son MD, PhD for their academic advice during this project. This Research was supported partly by the Intramural Research Program of the NIH, National Cancer Institute, Center for Cancer Research. This project has been funded in part with federal funds from the National Cancer Institute, National Institutes of Health, under contract HHSN26120080001E. The content of this publication does not necessarily reflect the views or policies of the Department of Health and Human Services, nor does the mention of trade names, commercial products, or organizations imply endorsement by the U.S. Government.

## Notes

**Conflict of interest:** The authors have no conflicts of interest to disclose.

